# The mitochondrial regulation of smooth muscle cell proliferation in type 2 diabetes

**DOI:** 10.1101/2023.02.15.528765

**Authors:** Olha M. Koval, Emily K. Nguyen, Dylan J. Mittauer, Karima Ait-Aissa, William Chinchankar, Lan Qian, Muniswamy Madesh, Dao-Fu Dai, Isabella M. Grumbach

**Author notes:** To whom correspondence should be addressed: Isabella Grumbach, MD, PhD, University of Iowa, Dept. of Internal Medicine, 169 Newton Road, Iowa City, IA 52246.

## Abstract

**Background:** Type 2 diabetes (T2D) is associated with a strongly increased risk for restenosis after angioplasty driven by proliferation of vascular smooth muscle cells (VSMCs). Here, we sought to determine whether and how mitochondrial dysfunction in T2D drives VSMC proliferation with a focus on ROS and intracellular [Ca^2+^] that both drive cell proliferation, occur in T2D and are regulated by mitochondrial activity.

**Methods:** Using a diet-induced mouse model of T2D, the inhibition of the mitochondrial Ca^2+^/calmodulin-dependent kinase II (mtCaMKII), a regulator of Ca^2+^ entry via the mitochondrial Ca^2+^ uniporter selectively in VSMCs, we performed in vivo phenotyping after mechanical injury and established the mechanisms of excessive proliferation in cultured VSMCs.

**Results:** In T2D, the inhibition of mtCaMKII reduced both neointima formation after mechanical injury and the proliferation of cultured VSMCs. VSMCs from T2D mice displayed accelerated proliferation, reduced mitochondrial Ca^2+^ entry and membrane potential with elevated baseline [Ca^2+^]_cyto_ compared to cells from normoglycemic mice. Accelerated proliferation after PDGF treatment was driven by activation of Erk1/2 and its upstream regulators. Hyperactivation of Erk1/2 was Ca^2+^-dependent rather than mitochondrial ROS-driven Ca^2+^-dependent and included the activation of CaMKII in the cytosol. The inhibition of mtCaMKII exaggerated the Ca^2+^ imbalance by lowering mitochondrial Ca^2+^ entry and increasing baseline [Ca^2+^]_cyto_, further enhancing baseline Erk1/2 activation. With inhibition of mtCaMKII, PDGF treatment had no additional effect on cell proliferation. Inhibition of activated CaMKII in the cytosol decreased excessive Erk1/2 activation and reduced VSMC proliferation.

**Conclusions:** Collectively, our results provide evidence for the molecular mechanisms of enhanced VSMC proliferation after mechanical injury by mitochondrial Ca^2+^ entry in T2D.

## INTRODUCTION

Vascular occlusive disease remains a critical cardiovascular health issue. In the US alone, it results in approximately 500,000 percutaneous coronary and 50,000 peripheral balloon angioplasties annually ^1^. Despite the use of contemporary drug-eluting stents during angioplasty, restenosis occurs in 3 to 20% of the treated arterial segments ^2^. In 2020, repeat procedures for in-stent restenosis represented over 10% of all angioplasties ^3^. One independent multivariate predictor of angiographic restenosis is type 2 diabetes mellitus (T2D) ^4,5^.

Restenosis is caused largely by neointimal hyperplasia, i.e., the proliferation of vascular smooth muscle cells (VSMCs) in the tunica media, and their subsequent migration to the intima. VSMCs that are derived from patients with T2D or treated with insulin and glucose proliferate at an abnormally high rate ^6,7^. Numerous cytosolic signaling pathways and downstream events have been implicated, including signaling via protein kinase C, the production of advanced glycation end products, the formation of reactive oxygen species (ROS) in response to ERK1/2/NFκB signaling, and the expression of PDK4 ^8–10^. Although altered mitochondrial function in the myocardium has also been reported in T2D ^11^, it has remained unclear whether and how it contributes to increased VSMC proliferation and restenosis after balloon angioplasty ^12^.

One means by which both cytosolic signaling and mitochondrial activity might contribute to cell proliferation is their control of cytosolic [Ca^2+^] ([Ca^2+^]_cyto_). For example, increases in [Ca^2+^]_cyto_ and Ca^2+^ transients are mechanisms by which growth factors promote cell proliferation ^13,14^. Indeed, in various cell types from patients with T2D or mouse models of diabetes ^15–17^ [Ca^2+^]_cyto_ is elevated, which has been attributed to both reduced Ca^2+^ export across the plasma membrane and impaired Ca^2+^ handling by the ER ^16,18–21^. We previously reported that both the proliferation ^22^ and migration ^23^ of VSMCs under normoglycemic conditions depend on mitochondrial Ca^2+^ uptake, in particular in G1/S phase when intracellular ATP demands are high. However, the extent to which T2D alters Ca^2+^ sequestration by mitochondria and whether this affects VSMC proliferation through retrograde signaling to cytosolic pathways in T2D is unknown.

The movement of Ca^2+^ into the mitochondrial matrix occurs through Mitochondrial Calcium Uniporter (MCU), which is a selective inward rectifying channel. Furthermore, this movement is driven by the highly negative inner mitochondrial membrane potential (ΔΨm). Importantly, the density of MCU channels in the inner mitochondrial membrane, as well as their subunit composition, vary greatly across tissue and cell types ^24^. Thus, the extent to which mitochondrial Ca^2+^ entry through MCU modulates cell physiology and human disease phenotypes is incompletely understood. With regards to diabetes, the data on mitochondrial Ca^2+^ sequestration from various rodent models are inconsistent and are currently limited to cardiac and skeletal muscle ^25–27^.

In this study, we tested whether enhanced neointima formation and VSMC proliferation in a diet-based mouse model of T2D can be reduced by manipulations that decrease mitochondrial Ca^2+^ entry. Specifically, we used the approach of inhibiting the Ca^2+^/calmodulin-dependent kinase II (CaMKII) in mitochondria by expression of the potent and highly selective inhibitor peptide CaMKIIN targeted to the mitochondrial matrix ^23,28,29^. Despite some disputation, mitochondrial CaMKII (mtCaMKII) is believed to promote mitochondrial Ca^2+^ entry by phosphorylation of the MCU at Ser^92 30–32^.

With this model, we establish the extent to which altered mitochondrial Ca^2+^ sequestration versus mitochondrial oxidative stress drive VSMC proliferation in T2D and link our finding to the retrograde activation of cytosolic growth pathways. Our data highlight the importance of mitochondrial Ca^2+^ handling in cell proliferation in T2D and suggest that it might be a particularly effective strategy for combatting neointimal hyperplasia in patients with T2D.

## RESULTS

### Neointimal hyperplasia in T2D mice is limited by inhibition of mitochondrial CaMKII

The Ca^2+^/calmodulin-dependent kinase CaMKII resides in mitochondria, where it is believed to activate MCU by phosphorylating Ser^92 33^. CaMKII is activated in the cytosol in various models of diabetes ^34,35^. As a first step toward ascertaining whether a genetic model in which targeting the CaMKII inhibitor peptide CaMKIIN to the mitochondrial matrix (mtCaMKIIN) ^29^ in VSMCs can provide insight into VSMC biology in T2D, we measured CaMKII oxidation and autophosphorylation in WT VSMCs (Figure S1A-C). Our results confirmed that acute hyperglycemia induces CaMKII activation in mitochondria. Also, adenovirus-mediated transduction of mtCaMKIIN blocked hyperglycemia-induced mitochondrial superoxide production in VSMCs treated with high glucose (Figure S1D), as well as their proliferation after treatment with insulin and high glucose (Figure S1E).

To test whether the mitoCaMKIIN reduces neointima formation *in vivo*, we placed mice expressing mtCaMKIIN selectively in smooth muscle cells^23^ and littermate controls on a high-fat diet (HFD, 60% fat) for 8 weeks and then injected them with a low dose of streptozotocin (STZ) to model the loss of pancreatic β-cells seen in humans with T2D. This model recapitulated the T2D phenotypes, including increased body weight (Figure S2A), impaired glucose tolerance and baseline blood glucose levels above 200 mg/dl, hypercholesteremia, hyperinsulinemia (Figure 1A-C). No significant differences were observed between the genotypes. The subsequent endothelial denudation of the common carotid artery using an epoxy bead-covered suture induced neointima formation in both genotypes^36,37^. However, the size of the neointima was significantly smaller in arteries from mice expressing mtCaMKIIN (Figure 1D-G, Figure S2B). Additional immunoblots of carotid artery tissue revealed that rates of apoptosis were similar in WT and mtCaMKIIN mice, suggesting that the observed differences in cell counts are due to decreased proliferation rather than to enhanced cell death (Figure S2C, D).

**Figure 1.**
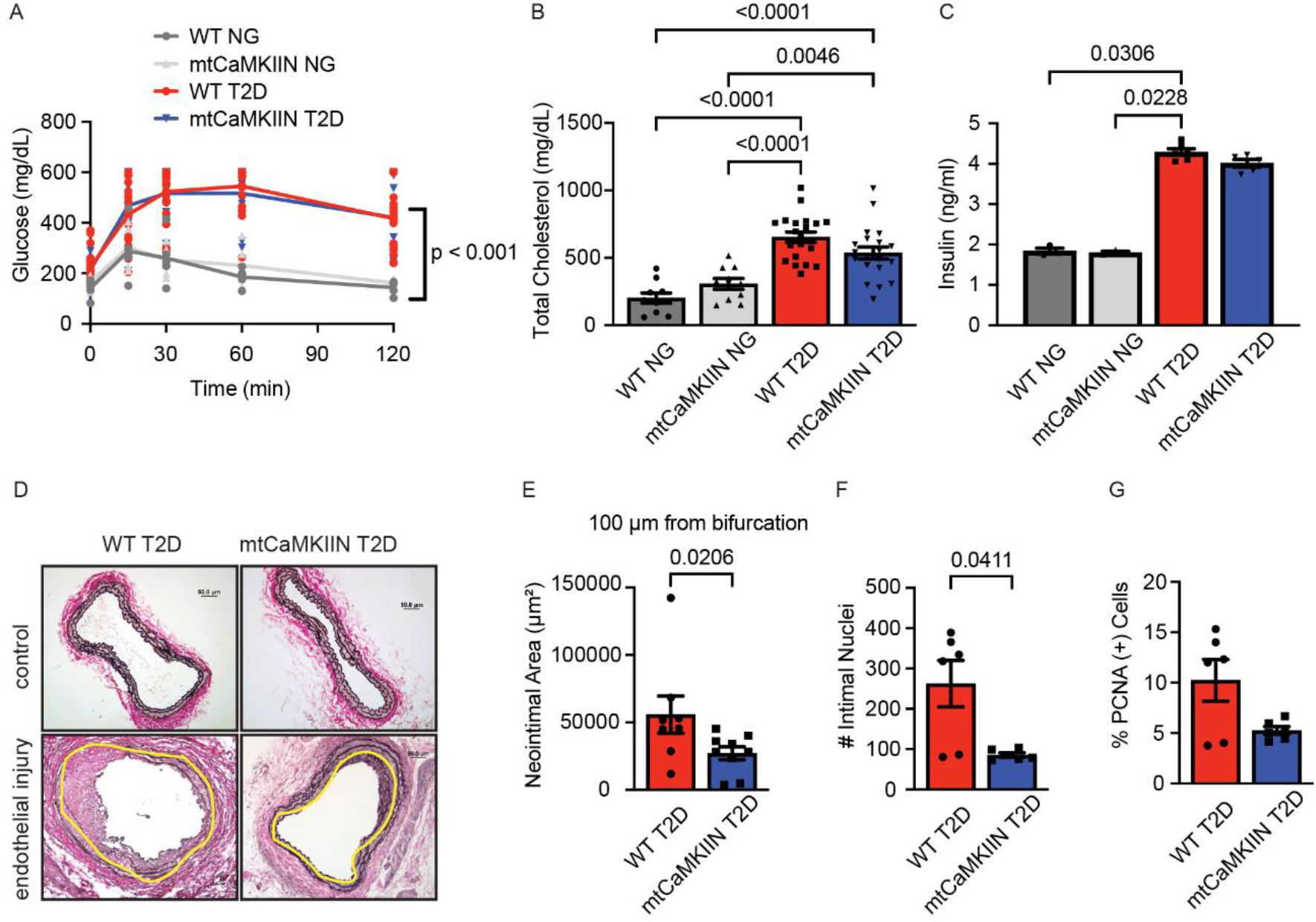
The mitochondrial CaMKII inhibitor mtCaMKIIN prevents neointimal hyperplasia after vascular injury. (A) Blood glucose levels over time after loading with glucose (2.0 g/kg i.v.), in normoglycemic mice (NG) and diabetic (T2D) mice of the wild-type (WT) and mtCaMKIIN genotypes. (B-C) Levels of (B) cholesterol and (C) insulin in serum from NG and T2D mice of the WT and mtCaMKIIN genotypes. (D) Verhoeff-Van Gieson staining of carotid arteries 21 days after endothelial denudation. Yellow line demarcates internal elastic lamina. (E) Neointimal area at 100 μm from bifurcation of the common carotid artery. (F) Number of nuclei and (G) percentage of PCNA-positive cells in the neointima, at same level as in (E). Analyses by two-way ANOVA (A), one-way ANOVA (B), Kruskal-Wallis test (C), and Whitney-Mann test (E-G).

### Enhanced proliferation of VSMCs isolated from T2D mice is dependent on mtCaMKII

To confirm that MCU inhibition blocks VSMC proliferation, we cultured aortic VSMCs from T2D and normoglycemic mice in which mtCaMKIIN was expressed in smooth muscle. VSMCs from T2D mice proliferated significantly faster than those from normoglycemic mice, in particular in the presence of the growth factor PDGF (Figure 2A). Notably, in the T2D mice the expression of mtCaMKIIN led to a significant reduction in cell proliferation. Proliferation was also strongly inhibited in VSMCs from T2D mice in which mtCaMKII activity had been impaired by adenoviral overexpression of the inhibitor peptide mtCaMKIIN (Figure 2B) or by treatment with the MCU inhibitor Ru265 (Figure 2C). Notably, the addition of the pharmacological MCU inhibitor Ru265 to VSMCs in which mtCaMKIIN was overexpressed did not affect cell proliferation, supporting that inhibition of CaMKII in mitochondria blocks proliferation through blockade of MCU and hence, of mitochondrial Ca^2+^ entry (Figure 2D).

**Figure 2.**
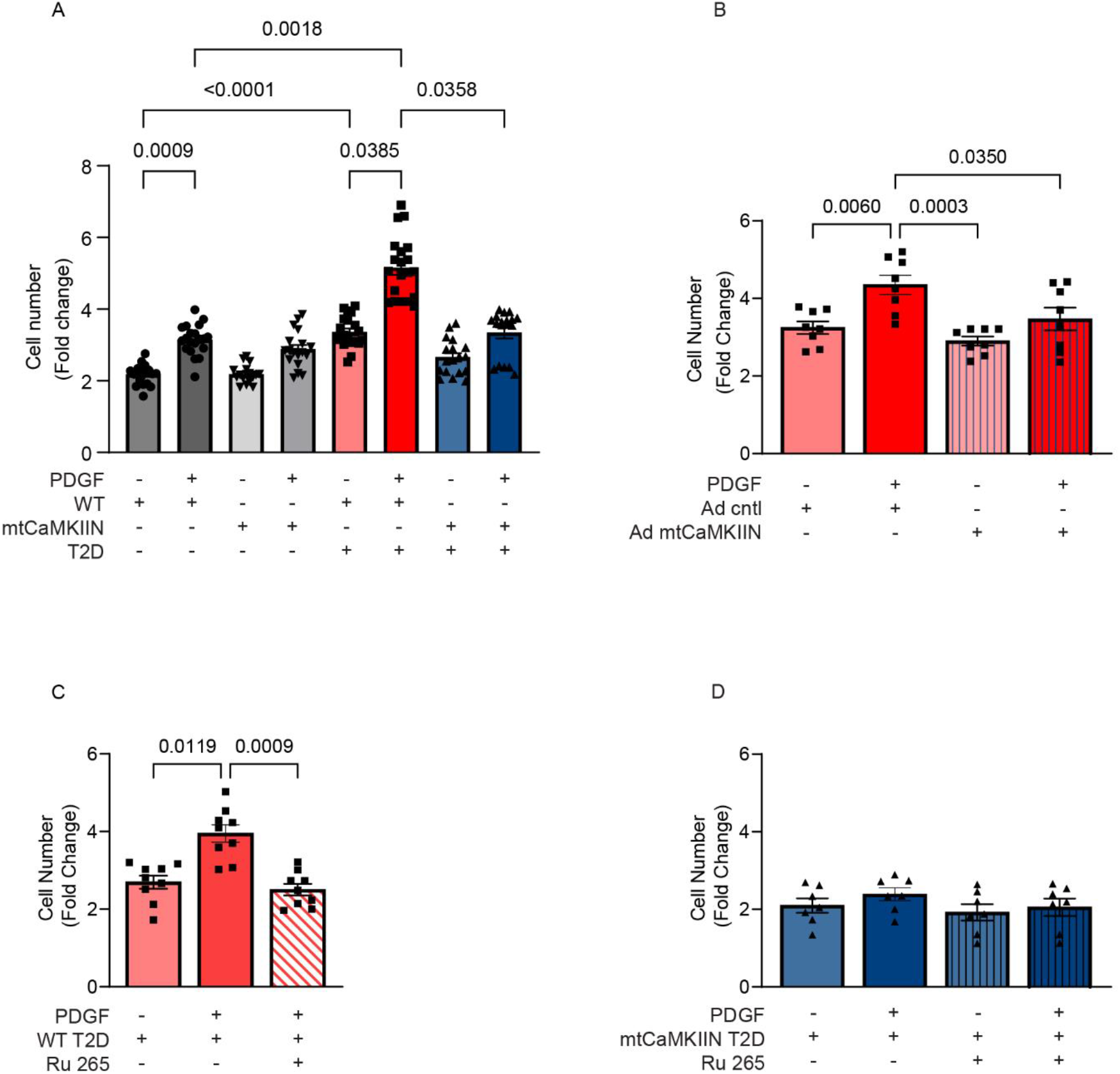
Inhibition of mitochondrial CaMKII reduces proliferation of VSMCs isolated from T2D mice. Numbers of VMSCs isolated from NG or T2D mice of the WT and mtCaMKIIN genetic backgrounds, counted after 72 hr in culture with or without PDGF (20 ng/ml). Data are expressed as fold change over levels at 0 hr. (A) VSMCs of the WT and mtCaMKIIN genetic backgrounds. (B) VSMCs isolated from T2D mice of the WT background and transduced with control or mtCaMKIIN adenovirus. (C) VSMCs isolated from T2D mice of the WT background, with or without administration of the MCU inhibitor Ru 265 (100 μM). (D) VSMCs isolated from T2D mice of the mtCaMKIIN background, with or without administration of the MCU inhibitor Ru 265 (100 μM). Analyses by Kruskal-Wallis test (A) and one-way ANOVA (C-E).

In mitochondrial preparations from VSMCs isolated from diabetic mice, MCU phosphorylation at Ser^92^ and CaMKII phosphorylation at Thr^287^ were detected (Figure 3A, B, C), indicating that MCU as well as mtCaMKII are activated in T2D. These findings would support increased mitochondrial Ca^2+^ entry. Thus, we imaged cytosolic Ca^2+^ clearance using Fura-2 following administration of boluses of 0.5 μM Ca^2+^ to permeabilized cells treated with thapsigargin (Figure 3D). Ca^2+^ clearance was enhanced in WT T2D VSMCs.

**Figure 3.**
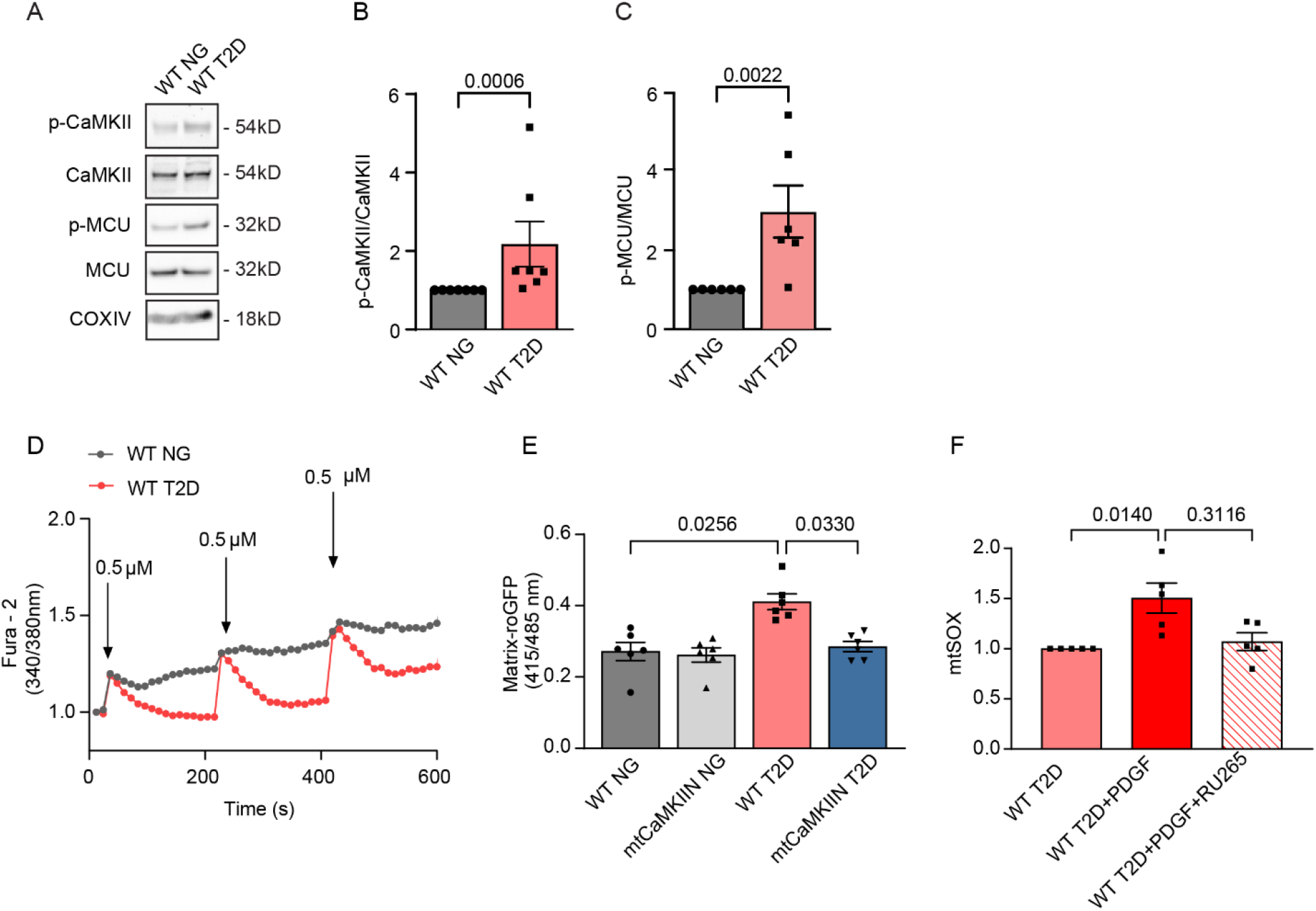
In VSMCs isolated from T2D mice, mtCaMKII is activated and drives mitochondrial ROS production. (A) Representative immunoblots assessing phosphorylation (i.e., activation) of CaMKII (at Thr^287^) and MCU A (at Ser^92^) in mitochondrial fractions from cultured VSMCs isolated from NG and T2D WT mice. (B, C) Quantification of (B) active CaMKII (p-Thr^287^), adjusted for CaMKII, and (C) active MCU A (p-Ser^92^), adjusted for MCU. (D) Cytosolic Ca^2+^ clearance as assessed by Fura-2, in response to 0.5 μM CaCl_2_ in permeabilized VSMCs from WT NG and T2D mice. (E) Mitochondrial ROS production as assessed by mito-roGFP in VSMCs from NG and T2D mice of the WT and mtCaMKIIN genotypes, induced by PDGF application (20 ng/ml). (F) Mitochondrial ROS production as assessed by mitoSOX in VSMCs from T2D after pretreatment with Ru265. Analyses by Mann-Whitney test in (B,C) and by Kruskal-Wallis test in (E, F).

In a previous study, we reported that mitochondrial ROS (mitoROS) production is dependent on Ca^2+^ entry ^39^. Here, in T2D, mitoROS production determined with the ratiometric indicator mitochondrial roGFP was elevated compared to normoglycemic conditions, and mtCaMKII inhibition reduced ROS production (Figure 3E). These findings were confirmed in mitoROS assays in which MCU was inhibited with Ru265 (Figure 3F).

### Inhibition of mitochondrial Ca^2+^ entry alters Ca^2+^ transients in response to PDGF

Next, we used mtPericam imaging to record [Ca^2+^] in the mitochondrial matrix in response to PDGF. The baseline [Ca^2+^]_mito_ was not significantly different in T2D versus normoglycemic VSMCs (Figure 4A). However, mitochondrial Ca^2+^ entry in response to PDGF was blunted in WT VSMCs from T2D mice, and it was reduced in the presence of mtCaMKIIN (Figure 4B-C) as also verified in HEK cells with mtCaMKIIN expression (Figure S3A).

**Figure 4.**
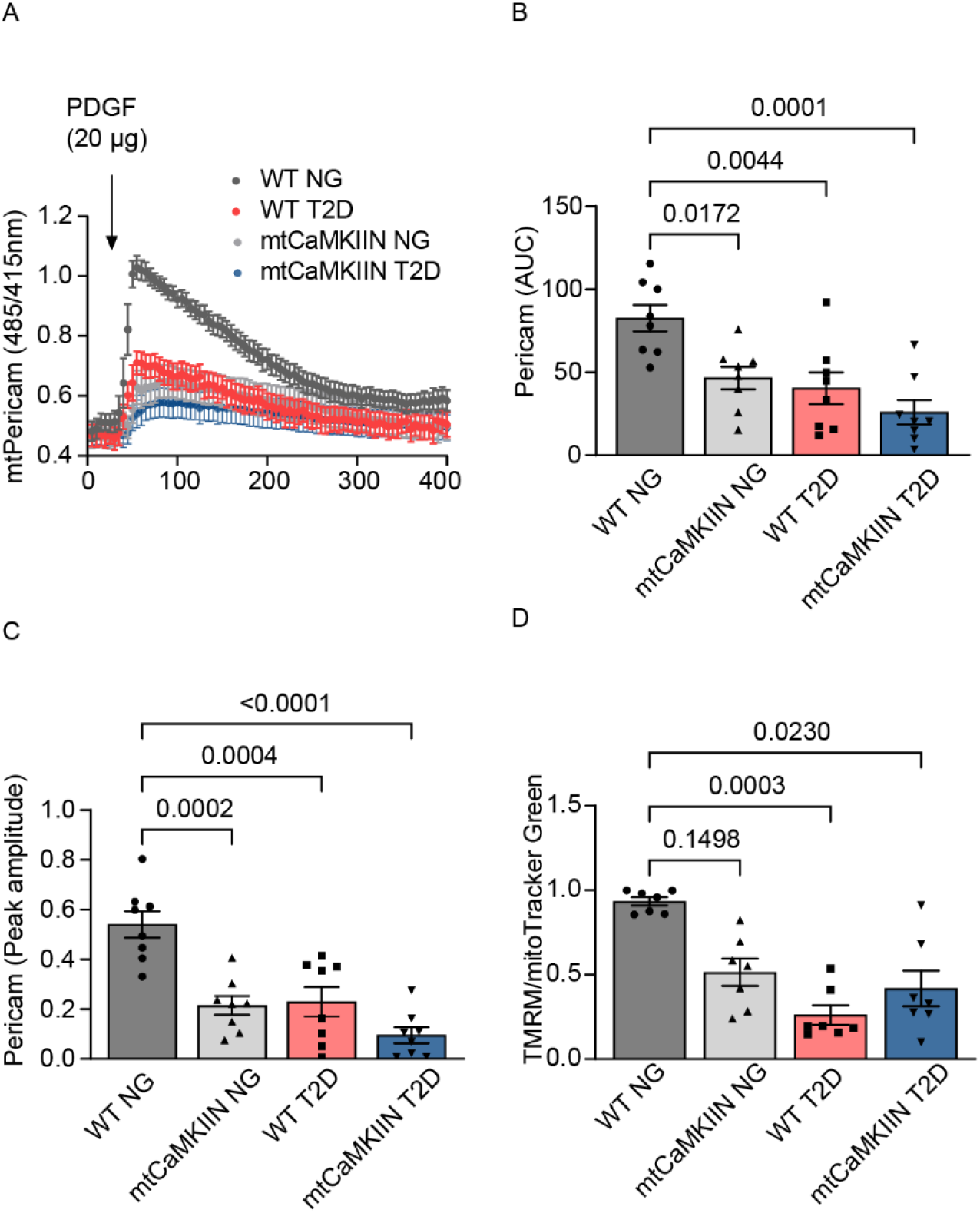
In VSMCs isolated from T2D mice, PDGF-induced mitochondrial Ca^2+^ transients are decreased. (A) Mitochondrial Ca^2+^ uptake as assessed by mtPericam imaging, in VSMCs from NG and T2D mice of the WT and mtCaMKIIN genotypes, induced by PDGF application (20ng/ml). (B, C) Quantification of (B) area under the curve (AUC) and (C) peak amplitude of mtPericam recordings from (A). (D) Mitochondrial membrane potential as assessed by TMRM imaging followed by normalization to the mitoTracker signal in VSMCs from NG and T2D mice of the WT and mtCaMKIIN genotypes. Analyses by Kruskal-Wallis test (B, C, D).

To provide a mechanism underlying decreased Ca^2+^ entry in intact, non-premetallized VSMCs from mice with T2D, we measured the mitochondrial membrane potential. VSMC mitochondria from diabetic mice were more depolarized than those from normoglycemic mice, providing some mechanistic underpinnings for the observed decreased Ca^2+^ entry. The mitochondrial membrane potential was not significantly influenced by the expression of mtCaMKIIN (Figure 4D). We ruled out that differences in the number of mitochondria per cell drive the variances in mitochondrial Ca^2+^ transients, we assessed the ratios of mitochondrial to nuclear DNAs for ND1/HK2 and Cox1/HK2. No significant differences were seen between cells from T2D mice with and without mtCaMKIIN expression (Figure S3B, C).

To establish how T2D and reduced mitochondrial Ca^2+^ entry affect [Ca^2+^]_cyto_, we measured baseline levels and cytosolic transients in response to PDGF. In VSMCs from WT T2D mice, the baseline [Ca^2+^]_cyto_ was higher than in normoglycemic mice of the same background (Figure 5A, B). Moreover, [Ca^2+^]_ER_ was lower in T2D, and inhibition of mtCaMKII had no additional effect (Figure 5C-E). Next, we tested whether T2D affects [Ca^2+^]_ER_ by altering the expression and activity of key cytosolic Ca^2+^ handling proteins, such as SERCA. Immunoblotting revealed that the levels of SERCA were decreased in T2D VSMCs, those of IP3R were increased and differences in NCX were not significant (Figure S4 A-D), supporting increased Ca^2+^ loss from the ER in T2D. Cytosolic Ca^2+^ transients triggered by PDGF were blunted in VSMCs from T2D mice (Figure 5F-H). Expression of mtCaMKIIN led to further increases in the baseline [Ca^2+^]_cyto_ and diminished the transients triggered by PDGF.

**Figure 5.**
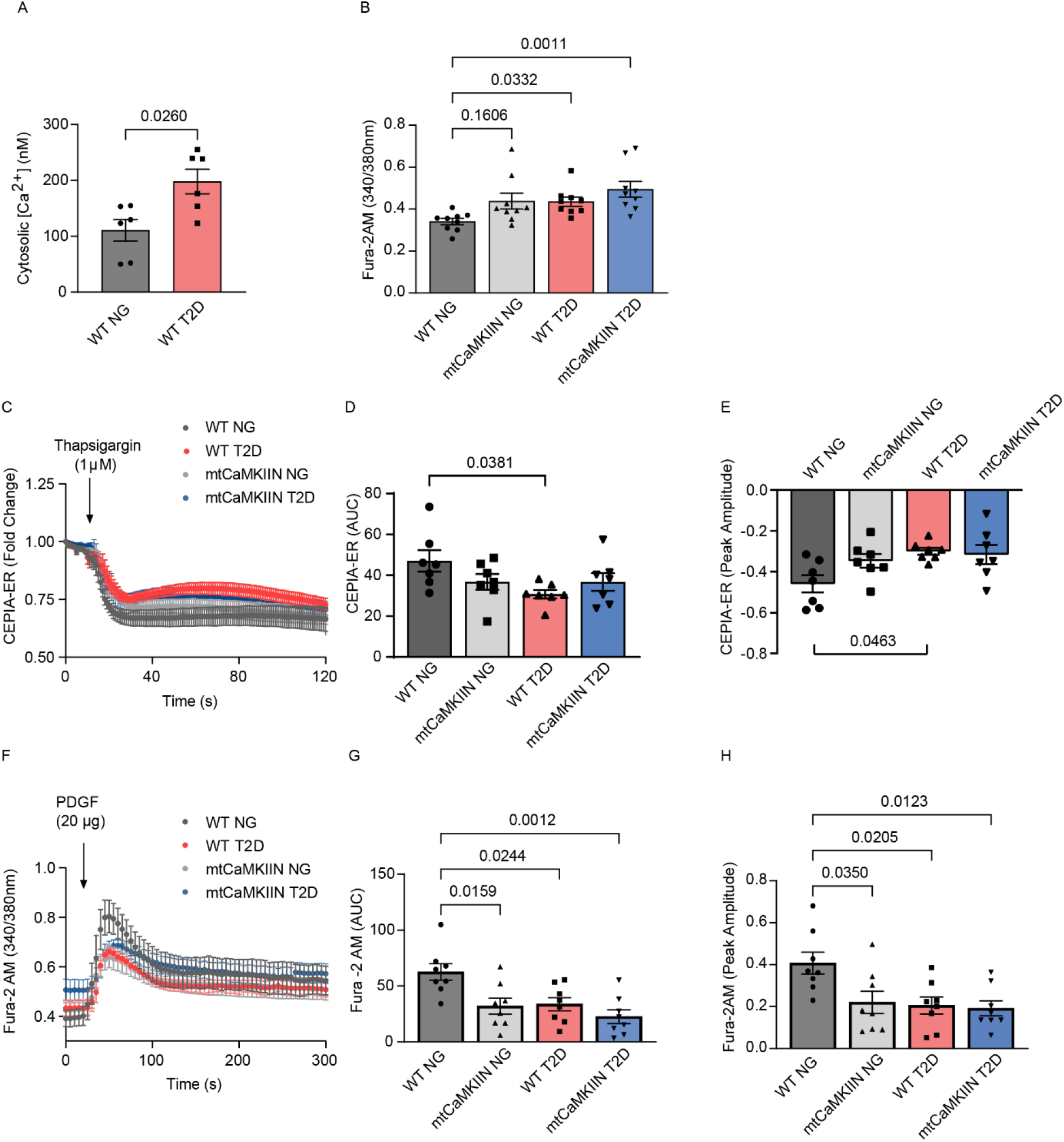
In VSMCs isolated from T2D mice, PDGF-induced cytosolic Ca2+ transients are decreased. (A) [Ca2+]_cyto_ in VSMCs from WT NG and T2D mice. (B) Baseline [Ca^2+^], as assessed by Fura-2AM imaging, in VSMCs from NG and T2D mice of the WT and mtCaMKIIN genotypes. (C, D, E) ER Ca^2+^-release induced by thapigargin (1μM). (D, E) Quantification of (D) area under the curve (AUC) and (E) peak amplitude of CEPIA-ER recordings from (C). (F) Cytosolic Ca^2+^ transients in response to PDGF. (G) Area under the curve (AUC) and (H) peak amplitude for Ca^2+^ transients, as in (F). Analyses by Mann-Whitney test (A) and Kruskal-Wallis test (B, D, F, G, H).

These data demonstrate that in T2D the baseline [Ca^2+^]_cyto_ is elevated. This is consistent with decreased levels of SERCA and increased expression of IP3R. The inhibition of mitochondrial Ca^2+^ entry by mtCaMKII inhibition enhances the cytosolic Ca^2+^ “overload” and further diminishes dynamic cytosolic Ca^2+^ transients after PDGF application without affecting [Ca^2+^]_ER_.

### Exaggerated proliferation of VSMCs isolated from T2D mice is driven by [Ca^2+^]_cyto_-regulated MAP kinase activation

Since cell proliferation is driven mainly by extramitochondrial events, i.e., cytosolic and nuclear signaling, we screened for differential activation of canonical growth pathways such as those involving MAP kinases. In initial experiments on normoglycemic and T2D VSMCs with expression of mtCaMKIIN, we did not detect differences in activation of the MAP kinases Akt and p38 (data not shown). However, in VSMCs from T2D mice, Erk1/2 phosphorylation was significantly higher than in samples from normoglycemic controls, and it increased further in the presence of mtCaMKIIN (Figure 6A). These results are consistent with previous reports that Erk1/2 activity can be enhanced by an increase in [Ca^2+^]_cyto_^40,41^. Upstream regulators of the Erk1/2 signaling pathway, such as c-Raf and MEK were activated in VSMCs isolated from T2D mice (Figure 6B).

**Figure 6.**
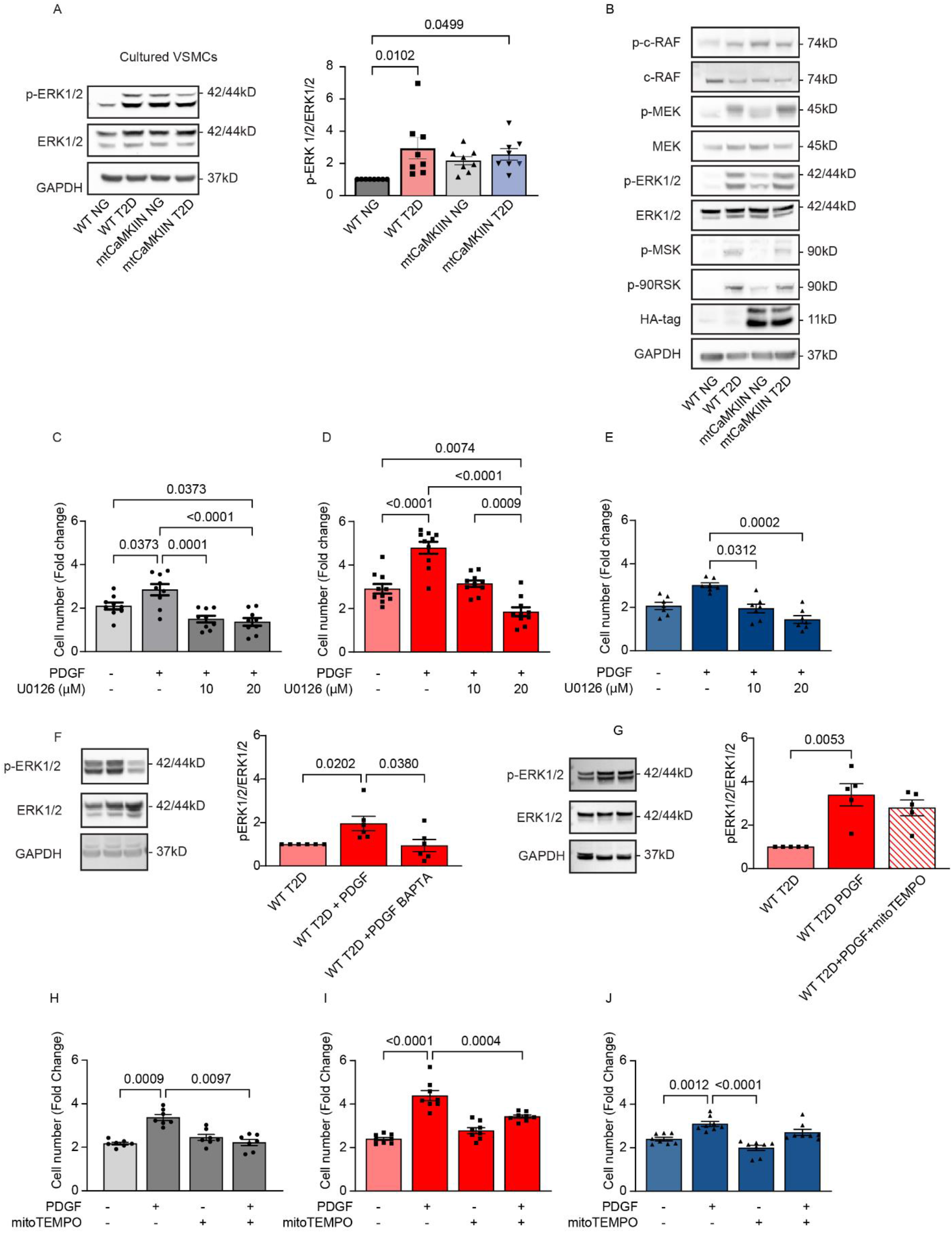
The enhanced proliferation of VSMCs isolated from T2D mice is caused by Ca^2+^-dependent Erk1/2 proliferation. (A) Representative immunoblots for phosphorylated (active) Erk1/2 and total Erk1/2 protein, and their quantification for cultured VSMCs isolated from NG and T2D WT and mtCaMKIIN mice. (B) Representative immunoblots for signaling pathway components upstream of Erk1/2 activation in cultured VSMCs from T2D mice of the WT and mtCaMKIIN genetic backgrounds. (C-E) Fold changes in cell number in (C) WT NG, (D) WT T2D and (E) mtCaMKIIN T2D VSMCs 72 hr after addition of PDGF and the Erk1/2 inhibitor U0126. (F) Representative immunoblot assessing phosphorylated (active) Erk-1/2 and (total) Erk1/2 protein levels and their quantification in cultured cells isolated from T2D mice of the WT genotype 15 min after addition of PDGF and preincubation with the Ca^2+^ chelator BAPTA (10 μM for 1 hr). (G) Representative immunoblot assessing phosphorylated (active) Erk-1/2 and (total) Erk1/2 protein levels and their quantification in cultured cells isolated from T2D mice of the WT genotype 15 min after addition of PDGF and preincubation with the ROS scavenger mitoTEMPO (10 μM for 1 hr). Fold changes in cell number in (H) WT NG, (I) WT T2D and (J) mtCaMKIIN T2D VSMCs 72 hr after addition of PDGF and the mitoTEMPO (10 μM). Analyses by Kruskal-Wallis test (A, C-E, H-G) and one-way ANOVA (F, J).

We then confirmed that activation of Erk1/2 contributes to accelerated proliferation in T2D. The concentration of the ERK1/2 inhibitor U0126 required to halt the proliferation of VSMCs was higher in WT T2D vs. WT normoglycemic mice, supporting the notion that Erk1/2 activity drives cell proliferation in T2D (Figure 6C, D). In VSMCs from T2D mice with mtCaMKII inhibition, U0126 had a reduced effect on proliferation (Figure 6E). We also ascertained that Erk1/2 activation depended directly on [Ca^2+^]_cyto_ by assessing Erk1/2 activation after pretreatment with the Ca^2+^ chelator BAPTA (Figure 6F). The activation of Erk1/2 by PDGF was abolished by pretreatment with BAPTA but not by pretreatment with mitoTEMPO (Figure 6G). Nonetheless, the addition of mitoTEMPO blocked the proliferation of VSMCs from WT T2D (Figure 6I) more effectively than from mtCaMKIIN T2D mice (Figure 6J), suggesting that excessive VSMC proliferation in T2D is dependent on both elevated mitoROS and altered Ca^2+^ handling but the effects mediated by Erk1/2 are mainly driven by [Ca^2+^].

### Erk1/2 hyperphosphorylation in T2D is driven by CaMKII activation in the cytosol

Since CaMKII in the cytosol is a Ca^2+^-dependent upstream regulator of Erk1/2, we tested its activating autophosphorylation at Thr^286^. As anticipated, autophosphorylation of CaMKII was enhanced at baseline in whole cell lysates VSMCs from T2D mice (Figure 7A). The expression of mtCaMKIIN further enhanced activation of cytosolic CaMKII consistent with an increase in [Ca^2+^]_cyto_ (Figure 7A). CaMKII co-immunoprecipitated with c-Raf, the upstream regulator of Erk1/2 (Figure 7B), indicating that CaMKII drives Erk1/2 activation. The addition of PDGF induced further association between these proteins in VSMCs from T2D WT mice. The expression of mtCaMKIIN enhanced the baseline association between c-Raf and CaMKII, but the treatment with PDGF did not increase the association beyond baseline levels (Figure 7C). Inhibition of cytosolic CaMKII by adenovirus-mediated expression of the untargeted inhibitor peptide CaMKIIN in the cytosol reduced baseline activation of CaMKII, MEK and Erk1/2, restored activation of the signaling pathway by PDGF (Figure 7C), and decreased VSMC proliferation relative to that in controls (Figure 7D).

**Figure 7.**
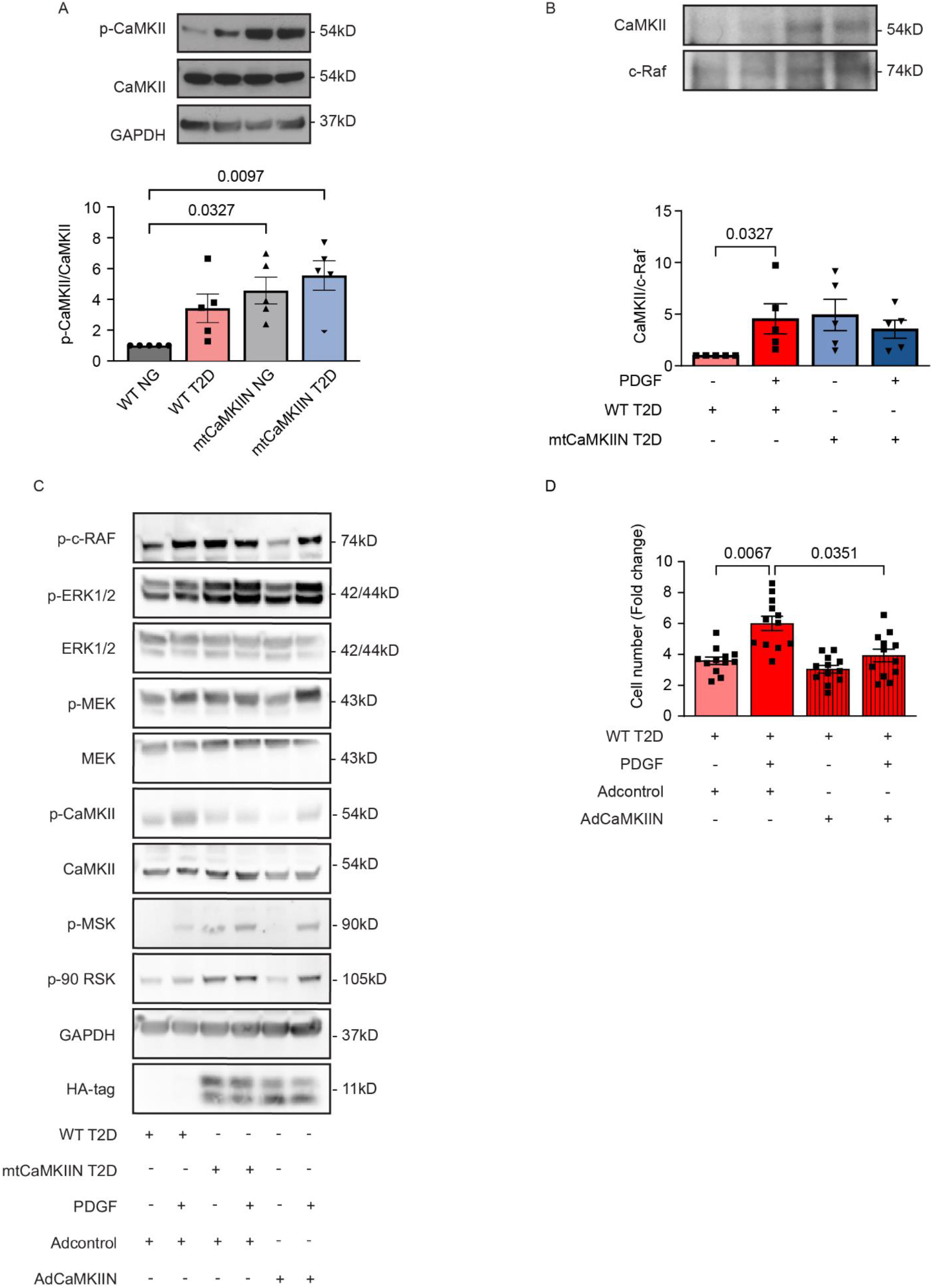
Activation of cytosolic CaMKII in T2D VSMCs mediates Erk1/2 activation. (A) Immunoblots for phosphorylated (active) CaMKII and total CaMKII protein in VSMCs isolated from NG and T2D mice of the WT and mtCaMKIIN genotypes. (B) Representative immunoblot (upper panel) and quantification (lower panel) of co-IP between CaMKII and c-Raf in VSMCs from diabetic (T2D) mice of the WT and mtCaMKIIN genotypes. (C) Representative immunoblots assessing upstream signaling components of the Erk1/2 signaling pathway in VSMCs from T2D mice of the WT and mtCaMKIIN genotypes. Cells were transduced with adenovirus expressing untargeted CaMKIIN for 72 hr prior to treatment with PDGF. (D) Number of VSMCs from T2D mice of the WT genotype following transduction with adenovirus expressing untargeted CaMKIIN or control virus for 72 hr prior to treatment with PDGF (20 ng/ml) and counted 72 hr later. Analyses by Kruskal-Wallis test (A, B, D).

## DISCUSSION

Our study demonstrates that inhibition of mitochondrial Ca^2+^ entry has a stronger antiproliferative effect in VSMCs from a model of T2D than in those from normoglycemic mice. We present four additional novel findings: First, in T2D, CaMKII is activated in VSMCs in mitochondria and thereby, controls mitochondrial Ca^2+^ entry, mitoROS production and proliferation. Secondly, inhibition of mtCaMKII in VSMC decreased neointima formation in vivo while having no effect on the development of T2D. Thirdly, enhanced proliferation of T2D VSMC was Erk1/2-dependent, was driven by altered [Ca^2+^]_cyto_ rather than by elevated mitoROS. Lastly, ablation of mtCaMKII further increased [Ca^2+^]_cyto_ and enhanced baseline Erk1/2 phosphorylation that could not be increased further by growth factor treatment and thereby, blocked proliferation.

Ca^2+^ uptake into the mitochondrial matrix through the MCU varies among different cells and tissues and is affected by extramitochondrial Ca^2+^ concentration, with low activity at resting and high capacity after cellular stimulation ^24,42^. Given the diversity in MCU expression and function in different tissues, the role of mitochondrial Ca^2+^ handling in human disease phenotypes is still under investigation. Evidence for altered expression of MCU subunits in cardiac and skeletal muscle in T2D has varied significantly with the particular models of diabetes studied ^25–27^. In contrast to numerous studies in cardiac myocytes and skeletal muscle, the extent to which mitochondrial Ca^2+^ handling affects VSMC function is less understood ^22,23^. Here, we present first evidence for MCU activation via mtCaMKII in VSMCs in T2D. Of note, diversity of MCU expression and function may also explain the reported discrepant effect of inhibition of mtCaMKII on mitochondrial Ca^2+^ in different tissues ^23,31–33,43–45^.

In our previous work, we demonstrated that mitochondrial Ca^2+^ entry drives VSMC migration under normoglycemic conditions using the same model system of inhibition by mtCaMKII ^23^. However, VSMC proliferation was not significantly inhibited by mtCaMKIIN expression in normoglycemia in contrast to our studies with MCU knockout, which we attributed to the lesser effect on mitochondrial Ca^2+^ entry by MCU inhibition with mtCaMKII inhibition versus MCU deletion ^22^. However, under altered mitochondrial Ca^2+^ handling in T2D with baseline mtCaMKII activation, its inhibition is sufficient to affect cytosolic proliferative signaling events.

Here, we also studied if T2D-induced Ca^2+^ signaling might be due to a dysregulation of either the cytosolic and ER Ca^2+^ homeostasis and/or the mitochondrial Ca^2+^ entry and whether blocking mitochondrial Ca^2+^ entry affects [Ca^2+^]_cyto_ or [Ca^2+^]_ER_. In human platelets and leukocytes of human subjects with T2D ^15,16,19^, [Ca^2+^]_cyto_ is elevated. This increase has been attributed to reduced export of Ca^2+^ across the plasma membrane and impaired Ca^2+^ handling by the ER ^16–21^, for example, in VSMCs by altered activity and expression of the K_ATP_ and the Cav1.2 channels ^46–49^. Our data demonstrate decreased [Ca^2+^]_ER_ in VSMC in T2D and congruent changes in protein expression of the IP3 receptor and of SERCA. Blocking the sequestration of [Ca^2+^] in mitochondria further elevated [Ca^2+^]_cyto_ in T2D, leading to increased preactivation of the Erk1/2 pathway. Surprisingly, Erk1/2 activation was driven by increased [Ca^2+^]_cyto_ rather than by elevated mitoROS.

Indeed, [Ca^2+^]_cyto_ is an accepted regulator of cell-cycle progression ^40^, and the mechanism is known to involve CaMKII ^50–53^ in the cytosol. Of note, increased activity of cytosolic CaMKII has been reported after hyperglycemia and in different models of diabetes in cardiac myocytes but not in cell proliferation ^34,35,54^. Although we have focused on the effects on cytosolic signaling events that promote proliferation, in future it will be important to investigate the extent to which mitochondrial metabolic activity (i.e., Ca^2+^-dependent production of ATP or intermediates of the TCA cycle) contributes to enhanced VSMC proliferation in T2D.

Our data provide the first evidence that MCU inhibition might be effective in combatting T2D-associated neointimal hyperplasia, a condition that continues to limit the use of percutaneous angioplasty in the T2D population. Recently, compounds with promise as therapeutics have emerged, for example the MCU inhibitor and Ruthenium derivative Ru265 and the novel smallmolecule inhibitors, such as MCU-i11 and MCU-i4 ^55^. However, significant neurologic side effects, including seizure activity, were reported after administration of Ru265 *in vivo* ^56^ and the *in vivo* efficacy of other inhibitors remains to be determined. As an attractive alternative, Crispr/Cas9 technology has been recently deployed to selectively ablate activated CaMKII ^57^. Given that mitochondrial Ca^2+^ entry is ubiquitous, effective use of any of these nascent tools for prevention of neointima formation would require the development of delivery strategies that are selective for VSMCs. Our data provide a strong rationale for testing inhibitors of mitochondrial Ca^2+^ entry for the treatment of vascular disease in T2D.

## METHODS

### Genetic mouse models

All animal procedures were approved by the University of Iowa Institutional Animal Care and Use Committee. This study was carried out in strict accordance with the recommendations in the Guide for the Care and Use of Laboratory Animals of the National Institutes of Health (NIH).

Mice of the C57BL/6 background that express tamoxifen-inducible Cre recombinase (driven by the smooth muscle myosin heavy chain, SMMHC, promoter) were obtained from Jackson Laboratories (#019079, denoted as “SM-Cre” mice). HA-tagged mtCaMKIIN mice were generated by cloning a cDNA that encodes an HA-tagged form of the CaMKII inhibitor peptide CaMKIIN (HA-CaMKIIN) fused to the mitochondria-targeting Cox8-palmitoylation sequence into a construct that contains the CX-1 promoter and a floxed eGFP sequence. Double-transgenic mice expressing mtCaMKIIN specifically in VSMCs (denoted as “mtCaMKIIN mice”) were generated by crossing SM-Cre^ERT2^ mice with HA-tagged mtCaMKIIN mice ^29^.. Eight-week-old, male SM-mtCaMKIIN mice were treated with a 5-day course of tamoxifen (20 mg/ml, i.p. injection, 5 days) to induce Cre recombination, and this regimen was repeated starting 15 days after the beginning of the first tamoxifen course.

SM-Cre mice treated with tamoxifen were used as controls for SM-mtCaMKIIN mice. Correct recombination and mtCaMKIIN expression were previously established (Nguyen et al). Because the SM-Cre transgene is on the Y-chromosome, only male mice were studied.

### Diet-induced type 2 diabetes

At 8-12 weeks of age, SM-mtCaMKIIN and SM-MCU^-/-^ mice and littermate control mice were fed a high-fat diet (HFD, 60% fat). After 8 weeks, streptozotocin (STZ) (75 mg/kg, 50 mg/kg) was administered by i.p. injections as two separate doses. Normoglycemic control mice were kept on normal chow. Induction of a T2D phenotype was verified by measuring body weight and glucose levels in blood from the tail vein. Two weeks after the STZ injection, HFD mice and normoglycemic controls were subjected to a 6-hr fast followed by a glucose tolerance test. Specifically, basal glucose levels were measured in blood from the tail vein. Subsequently the mice were i.p. injected with 20% glucose in saline (2.0 g/kg based on lean mass) and blood glucose levels were measured at 15, 30, 60 and 120 min thereafter.

### Serum Assays

Blood plasma samples were collected by tail-tip incision and centrifugation at 2500 rpm. Plasma insulin concentrations were measured using a mouse insulin ELISA kit (CrystalChem, 90080), and plasma cholesterol levels using a colorimetric cholesterol quantitation kit (Sigma-Aldrich, MAK043-1KT), according to the manufacturers’ protocols.

### Carotid Injury

Vascular injury was modeled in 10-week-old SM-mtCaMKIIN mice and littermates by endothelial denudation of the left common carotid artery using a resin bead-coated suture. Twenty-eight days after the denudation procedure, all animals were anesthetized. Transcardiac perfusion was performed with 10 ml PBS, after which fixation at physiological pressure was performed by replacing PBS with 10 ml of 4% paraformaldehyde (PFA). The injured left carotid arteries and their contralateral controls were excised, placed in tissue-embedding molds, stored in 4% PFA for 2 days, and embedded in paraffin.

For morphometric analysis of the arteries, 5 μm cross sections were obtained at 50 μm intervals, beginning at the carotid bifurcation site and extending up to 500 μm proximally. Sections were treated with Verhoeff-van Gieson stain and imaged using a Nikon Eclipse TS100 microscope. The areas of the vessel wall, lumen, and intima were determined using tracings of the circumferences of the external elastic lamina (EEL), internal elastic lamina (IEL), and lumen, using NIH Image J. The areas were calculated as follows: that of the vessel wall by subtracting the luminal area from the area defined by the EEL; and that of the neointima by subtracting the luminal area from the area defined by the IEL.

### Cell culture

Primary aortic VSMCs were isolated from mtCaMKIIN, and control mice by enzymatic digestion. The aortas were incubated in elastase (to remove adventitia) and cut into rings (about 1 mm in length) that were subsequently incubated in 2 mg/ml collagenase type II (Worthington Biochemical Corporation) for 2 hr. The digested pieces were plated in DMEM with 450 mg/dl glucose, 1% penicillin/streptomycin, MEM nonessential amino acids, MEM Vitamin and 8 mmol/L HEPES) supplemented with 20% fetal bovine serum (FBS) and 0.1% fungizone. The cells were cultured in DMEM with 10% FBS at 37°C in a humidified incubator (95% air and 5% CO_2_). Cells were routinely tested for mycoplasma contamination.

### Adenoviral transduction

Adenoviruses expressing mtCaMKIIN (Ad-mtCaMKIIN), untargeted CaMKIIN (Ad-CaMKIIN), or mito-Pericam (Ad-mtPericam), as well as empty vector (Ad-control), were generated by the Viral Vector Core Facility at the University of Iowa.

### Detection of mitochondrial ROS

Ro-GFP (reduced-oxidized GPP) plasmid targeted to mitochondrial matrix was purchased from Addgene (#49437) and delivered into cells by electroporation. The redox-sensitive protein excitations were induced at 400 and 484 nm, with emission measured at 525 nm.

Cells were loaded with mitoSOX red (5 μM; a dihydroethidium derivative, Invitrogen, M36008) and mitoTracker Deep Red (100 nM; fluorescent mitochondrial stain, Life Technology, M22426) for 20 min at 37°C. In some experiments, VSMCs had been transduced with empty adenovirus or mtCaMKIIN for 48-72 hr prior to this loading. After 20 min in the respective dye, the cells were washed with HBSS and imaged using a Zeiss 510 confocal microscope. Analysis was performed by tracing a region of interest around the cell using NIH Image J, and the mean fluorescence intensity of mitoSOX was normalized to the mitoTracker signal. Data are presented as relative fluorescence units.

### Measurement of cytosolic Ca^2+^ transients

Cells were loaded with Fura-2 acetoxymethyl ester (Fura-2AM) by incubation with 2 μM Fura-2AM in HBSS for 20 min at room temperature, then washed twice and incubated at 37°C for 5 min for de-esterification. Cells were excited alternatively at 340 and 380 nm. Intensity of the fluorescence signal was acquired at 510 nm. Real-time shifts in the Fura-2AM fluorescence ratio were recorded before, during, and after acute addition of PDGF (20 ng/ml) using a Nikon Eclipse Ti2 inverted light microscope. Peak amplitude was calculated by subtracting the baseline fluorescence ratio from the peak fluorescence ratio. The area under the curve (AUC) was determined using GraphPad Prism and normalized by subtracting the AUC at baseline. Summary data represent the average differences in the basal and peak increases in [Ca^2+^] cyto. The free [Ca^2+^] cyto was calculated from Fura-2AM fluorescence using the equation [Ca^2+^]_cyto_ = (R – R_min_) / (R_max_ – R) · K_d_ · β, where R is the basal ratio of fluorescence at 340 nm to 380 nm, K_d_ is the Fura-2AM dissociation constant (145 nM), and β is the difference in Fura fluorescence intensity in Ca^2+^-free vs. Ca^2+^-saturated media, according to the method described by Favaro et al. (2019).

### Measurement of mitochondrial Ca^2+^ transients

Ratiometric Ca^2+^ measurements in mitochondria were performed using adenovirus-delivered mtPericam (Ad-mtPericam), a fluorescent Ca^2+^ indicator protein with a COX IIIV targeting sequence. VSMCs were infected with Ad-mtPericam 48 hr prior to analysis. Ratiometric fluorescent imaging of mtPericam via Nikon Eclipse Ti2 microscope was used to determine the intensity of the fluorescence signal, with excitation at 415 nm and 480 nm, with emission at 510 nm. Peak amplitude and AUC were determined as described for Fura-2 AM.

### Measurement of ER Ca^2+^

Cepia ER plasmid was purchased from Addgene (#58215). The CEPIA1er protein was excited at 543 nm, and emitted fluorescence was measured at wavelengths of 580 nm.

VSCMs were transfected in a Nucleofector I device (Lonza) using the Basic Nucleofector Kit for Primary Mammalian Smooth Muscle Cells (VPI-1004, Lonza) and following the manufacturer’s protocol. Six hundred thousand cells were transfected in the presence of 5-μg of plasmid DNA, plated onto 35-mm glass bottom microwell dishes (MatTek Corporation), and grown for 48 hr before experiments were performed.

### Measurement of mitochondrial membrane potential

The mitochondrial membrane potential was measured using the tetramethylrhodamine methyl ester (TMRM, 50 nM, Life technology, T668) and Mitotracker Green (100nM, Thermo Fisher Scientific, M7514) fluorescent probes according to the manufacturers’ protocols. Images were taken at baseline and 15 min after FCCP treatment (5uM, Sigma, C2920), using an LSM 510 confocal microscope at a magnification of 40x (Carl Zeiss, San Diego, CA). Image analysis was performed using NIH ImageJ. All images were taken at the same time and using the same imaging settings. Data are presented as fold change over control with referencing to the completely depolarized state.

### Cell lysis and fractionation

Whole cells were lysed in RIPA buffer (20 mM Tris, 150 mM NaCl, 5 mM EDTA, 5 mM EGTA, 1% Triton X-100, 0.5% deoxycholate, 0.1% SDS, pH 7.4) supplemented with both protease inhibitors (Mini complete, Roche) and phosphatase inhibitors (PhosSTOP, Roche). Lysates were sonicated and debris was pelleted by centrifugation at 10,000 × g for 10 min at 4°C. Mitochondrial fractions were prepared in MSEM buffer (5 mM MOPS, 70 mM sucrose, 2 mM EGTA, 220 mM Mannitol, pH 7.5 with protease inhibitors), with homogenization performed in cold MSEM buffer in a Potter-Elvehjem glass Teflon homogenizer (50 strokes). Nuclei and cell debris were pelleted by centrifugation at 600 × g for 5 min at 4°C. Mitochondria were separated from the cytosolic fraction by centrifugation at 8000 × g for 10 min at 4°C. Protein concentrations were determined using the Pierce™ BCA protein assay (Thermo Scientific).

### Immunoblotting and immunoprecipitation

For immunoprecipitation assay, cells were lysed in Pierce IP Lysis Buffer (Thermoscientific, 87788). For each condition, 300 μg of proteins were used and incubated with primary antibody overnight at 4°C degrees, and then with magnetic Dynabeads Protein G (Invitrogen, 58096) for 1 hr at room temperature. Dynabeads with precipitated proteins were washed 3 times with DynaMag-2 (Invitrogen) and eluted with LDS buffer (Invitrogen, 58011).

Equivalent amounts of 5-15 μg of protein for gel loading (cell lysates, mitochondrial/cytoplasmic fractions) were separated by SDS/PAGE on 4-20% Tris/glycine precast gels (BioRad) and transferred to PVDF membranes (BioRad). Membranes were blocked in 5% BSA and incubated overnight at 4°C with primary antibodies. Blots were washed 3 times for 10 min with 0.05% Tween-20 in TBS, incubated for 1 hr at room temperature with the respective secondary antibodies, and then washed again. The blots were then developed using ECL chemiluminescent substrate (ThermoScientific 34580) according to the manufacturer’s instructions.

### Immunohistochemical staining

Human artery paraffin blocks were sectioned to 5 μm thickness. 1X PBS (0.145 M NaCl, 0.0027 M KCl, 0.0081 M Na_2_HPO_4_, 0.0015 M KH_2_PO_4_, pH 7.4) was used as washing buffer and PBS-based incubation buffer was supplemented with 1% bovine serum albumin, 1% normal donkey serum, and 0.3% Triton® X-100. Primary MICU1 antibody (1:150) and smooth muscle actin antibody (1:500) were applied overnight at 4°C. For development of the diaminobenzidine (DAB) signal, the ImmPRES S® Duet Double Staining Polymer Kit (vectorlabs.com, MP-7714) was used, and this was followed by counterstaining with hematoxylin.

### Proliferation Assays

VSMCs were cultured in 12-well plates at 5,000 cells per well in DMEM containing 10% FBS. At 24 hr after plating they were treated with PDGF (20 ng/ml), and at 72 hr after they were trypsinized and triplicate samples were counted using a Beckman Coulter Z1 cell counter.

### Ca^2+^ retention assay

Calcium Green-5N was used to monitor cytosolic Ca^2+^ in permeabilized VSMCs. Signal was recorded in 96-well plates. The total volume of the reaction in each well was 100 μl, including 50 μL of cell suspension (2 000 000) and 50 μl of respiration buffer (100 mM K aspartate, 20 mM KCl, 10 mM Hepes, 5 mM glutamate, 5 mM malate, and 5 mM succinate (pH 7.3)) supplemented with 5 μM thapsigargin, 0.005% digitonin, and 1 μM Calcium Green-5N (Invitrogen). Fluorescence was monitored at 485 nm excitation and at 535 nm emission. Following measurement of baseline Ca^2+^ levels, CaCl_2_ (10 μM Ca^2+^ at 30°C) was added 5 times every 4 min until Ca^2+^ uptake ceased.

### qPCR to determine mitochondrial DNA copy number

Genomic DNA was isolated from approximately 500,000 VSMCs using the DNeasy Blood & Tissue Extraction Kit (Qiaqen). DNA samples were treated with RNase and subsequently quantified using the Power SYBR Green Master Mix Real-time PCR Kit, using 100 ng genomic DNA per reaction (Thermofisher, 4367659). The mitochondrial DNA copy number was calculated using the 2^−ΔCt^ formula, where Δ = Ct_mito_ – Ct_nuclear_, for selected mitochondrial (ND1, COX1) and nuclear (HK2, NDUFV1) genes.

### Statistical Analyses

Data are expressed as mean ± SEM and were analyzed using the GraphPad Prism 9.0 software. All data sets were analyzed for normality and equal variance. The Kruskal-Wallis test with Dunn’s post hoc test was used to assess data sets where normal distribution could not be assumed. Student’s T-test and one-way ANOVA followed by Tukey’s multiple comparison test were used for data sets with normal distributions. Two-way ANOVA followed by Tukey’s multiple comparison test was used for grouped data sets. A p-value of <0.05 was considered significant.

## Autor contributions

O.K. - designed research studies, conducted experiments, acquired data, analyzed data, wrote the manuscript.

E.N. - conducted experiments, acquired data, analyzed data.

D.M. - conducted experiments, analyzed data.

K.A-A.- conducted experiments, analyzed data.

W.C. - analyzed data.

M.M. - provided reagents, wrote the manuscript.

D-F.D. - conducted experiments, acquired data, analyzed data.

I.G. - designed research studies, provided reagents, provided funding, wrote the manuscript.

## Acknowledgements

The authors thank Dr. Christine Blaumueller of the Scientific Editing and Research Communication Core at the University of Iowa Carver College of Medicine for critical reading of the manuscript.

## KEY RESOURCES TABLE

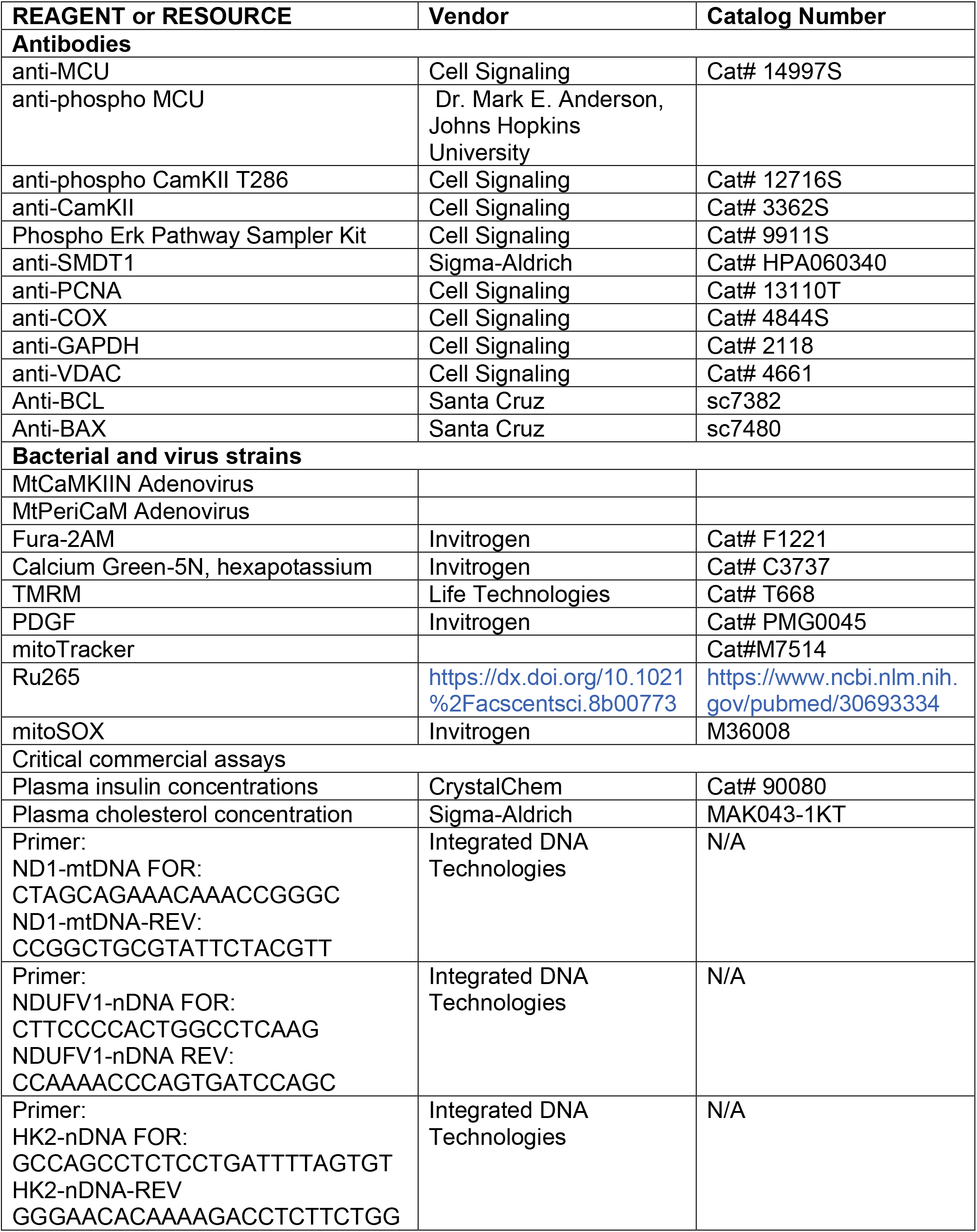

